# Cardiomyopathic mutations of lamin A perturb mutual interactions of lamin, nuclear membrane, and chromatin leading to LLPS

**DOI:** 10.1101/2024.10.07.616979

**Authors:** Subhradip Nath, Shuvadip Dutta, Shreyasi Dey Sarkar, Duhita Sengupta, Mithun K. Mitra, Kaushik Sengupta

## Abstract

Lamins are nuclear intermediate filaments constituting the nuclear lamina which maintains the structural integrity of the nucleus and play a key role in the spatial organisation of the genome. Mutations in the lamin protein have been associated with diverse diseases collectively known as laminopathies. In this study, we focused on two lamin A mutants - E161K and K97E - associated with dilated cardiomyopathy (DCM). Through confocal imaging, we established that these mutations cause large scale disruption of the peripheral lamin and consequent heterochromatin organisation, along with the formation of lamin aggregates inside the nucleoplasm. Using coarse-grained polymer simulations, we uncovered the role of lamin-lamin, lamin-membrane and lamin-chromatin interactions in maintaining wild-type lamin and chromatin organisation and showed that disruptions in these interactions can reproduce the experimental observations in the lamin mutants. These predictions were verified using 3D-FISH experiments to quantify the reorganisation of chromosome territories in these mutants. Using advanced imaging methods, we characterised the dynamical properties of the lamin aggregates in the mutants to show for the first time a liquid-like state of the lamin aggregates through a liquid-liquid phase separation. The altered lamin and chromatin interactions in the mutants thus manifest as liquid-like aggregates in the nucleoplasm leading to disruption of the spatial organisation of the genome in these laminopathy-associated mutants.

## I. INTRODUCTION

The spatio-temporal organization and dynamics of the human genome crucially coordinate the gene expression program inside cells thereby governing the basic cellular functions - proliferation, differentiation, and tissue homeostasis [1]. A key mediator affecting the positioning of the chromosomes inside the nucleus are lamin proteins which constitute the nuclear lamina (NL) [2–4], a 10-30nm thick filamentous network underlying the double-membrane bound nuclear envelope [5]. Nuclear lamins can mediate this interaction with chromatin either directly via its tail domain [6–8] or by binding to histone proteins [9, 10] or other chromatin-binding proteins such as emerin, MAN1 [11], lamin B receptor (LBR) [12–14], LAP2β [15], BAF [16] and HP1 [17]. Lamins are type-V intermediate filament proteins and are categorized into either A-type or B-type lamins. A-type lamins are encoded by *LMNA* gene and constitute lamin A, splice variant lamin C and lamin AΔ10 [18]. Likewise, lamin B1 and B2 account for the B-type lamins and are expressed from *LMNB1* and *LMNB2* genes respectively [19, 20]. A-type lamins are expressed minimally in pluripotent stem cells and during early embryogenesis but are noticeably upregulated with differentiation [21, 22] whereas B-type lamins are constitutively expressed in all cell types. Furthermore, B-type lamins are widely distributed along the nuclear periphery whereas A-type lamins are found in both the nuclear periphery and nucleoplasm [20]. Depletion of either A or B-type lamins has been associated with mis-localization of chromosome territories [23–28]. Genome-wide mapping methods have shown the association of the peripheral AT-rich chromatin domains, referred to as lamina associated domains (LADs), with the nuclear lamina [1, 29–31]. LADs vary between hundreds of kilobases to megabases in length and characterized by the preponderance of H3K9me2, H3K9me3 and H3K27me3 modifications [1, 32, 33]. Although LADs have predominantly been analysed with respect to lamin B1 proximity [29, 34, 35], the role of A-type lamins in determining chromosome organisation through interactions with chromatin remains unclear. The potential role of A-type lamins has been shown experimentally where LAD tethering in differentiated cells as well as chromatin dynamics is perturbed when A-type lamins are depleted [32, 36]. The interactions of lamin proteins with the LADs plays a crucial role in the organisation of the chromatin polymer. Hi-C experiments have revealed the compartmentalisation of chromatin into distinct megabase-scale regions - gene-rich and transcriptionally active A-type compartments and gene-poor and transcriptionally silent B-type compartments [37]. The peripheral organisation of heterochromatic B-type domains can then be understood in terms of the interactions of chromatin with the nuclear lamina [38].

The role of polymer physics in understanding cellular organization has been highlighted through studies of chromatin organization [39–42], interactions of chromatin with various proteins [43–45], and membraneless bio-molecular condensates at various length scales [46– 49]. These studies utilize polymer physics concepts to elucidate the mechanisms of phase separation and mechanical properties of bio-polymers within the crowded cellular environment. Several simulation-based studies have explored polymer modeling of chromatin along with the nuclear lamina, focusing on how chromatin interacts with lamin proteins and the implications for nuclear organization [50–54]. Earlier study on Lamin-associated chromatin model for chromosome organization presents a minimal polymer model that quantifies domain lengths and successfully reproduces the formation of lamin-associated domains (LADs), highlighting the significance of LADs in genome organization [55]. Heterogeneous interaction between different compartments of chromatin and lamin leads to either peripheral organisation of heterochromatin in a conventional nuclei or a central chunk of heterochromatin in inverted nuclei [56, 57]. Computational models have shown that interactions with the lamina can compact chromatin, with the degree of compaction proportional to the number of contact points [58]. Models also predict that LAD detachment from the lamina coincides with local decompaction of chromatin [58]. Systematic studies on interplay between chromatin-lamina interactions, intra-chromatin interactions, and hydration have provided insights into the chromatin organization [54]. Computational models are emerging to predict chromatin behavior at the lamina, offering insights into normal and disease states. Polymer models capture the stochastic nature of LADs and explains the reorganization of heterochromatin and euchromatin in senescent cells [59, 60]. Alteration of radial positioning of the LADs was found in lamin A mutants linked to a progeroid syndrome which aligned with the prediction from 3D genome modeling data [61–63]. Nearly 500 *LMNA* mutations have been linked to 14 different human diseases, collectively known as laminopathies, characterised by alteration of nuclear stiffness and misshapen/fragile nuclei. Laminopathies include dilated cardiomyopathy (DCM), Emery-Dreifuss muscular dystrophy (AD-EDMD), Hutchinson-Gilford progeria syndrome (HGPS), atypical Werner’s syndrome (WS), restricted dermopathy (RD), and mandibuloacral dysplasia (MAD) [18]. Dilated cardiomyopathy (DCM) is the most prevalent cardiac condition reported in cardiovascular clinics across the world and is characterized by ventricular dilation with impaired systolic functioning resulting in sudden cardiac arrest and death in patients [64]. Around 200 *LMNA* mutations have been associated with DCM to date accounting for 5-10% of DCM patients [65–67]. In case of laminopathies severe disorganization of chromatin in the 3D nuclear space were reported [68, 69]. Pathogenesis in muscular dystrophy and dilated cardiomyopathy caused by mutations in lamin A was found to be linked with altered heterochromatin distribution [70] and transcriptional landscape [71, 72].

The mutations K97E and E161K addressed in this study have been independently documented in cohorts of DCM patients from Italy, Spain, Germany, United States, Finland, Ireland, and Korea and cause severe phenotypes that lead to abrupt cardiac arrest and death [73–76]. Epigenetic changes have been implicated in several studies as one of the primary underlying causes of cardiovascular disorders [77–79]. It has been shown that the mutation E161K modifies the gene expression profile in the human cardiomyopathic heart [80]. A recent report also showed the modification of epigenetic landscape and its effect on the transcriptome caused by lamin A K97E [18].

In this study, we have focused on lamin A-chromatin interaction and dynamics with respect to the ectopic expression two DCM-associated mutants of lamin A, K97E and E161K, to delineate how the mutations in the rod domain of lamin A influence the lamin-chromatin interaction and consequent genome organisation in these mutants. Using confocal imaging and polymer modelling, we investigated how lamin-lamin and lamin chromatin interactions can lead to large scale changes in both the lamin and chromatin organisation. We have established for the very first time the formation of liquid-like phase separated condensates of lamin in the mutant nuclei, consistent with a corresponding shift in the lamin-lamin, lamin-membrane and lamin-chromatin interactions.

## II. RESULTS

### A. Lamin A mutants K97E and E161K show formation of lamin aggregates and altered heterochromatin organisation

The mutants E161K and K97E map in the rod 1B domain of lamin A as shown in Fig. 1 A. We investigated the effects of these two mutations on the global lamin and chromatin organization as compared to the WT. We transfected mouse myoblast cell line C2C12 with the EGFP-tagged lamin A and its mutants. Mutant transfected C2C12 cells are known to serve as a good model system for the human cardiomyocytes which are adversely affected in DCM [81]. We observed a diminishing peripheral lamina in E161K which completely disappeared in K97E concomitant with the formation of dense aggregates inside the nucleoplasm (Fig. 1 B) while the WT lamina was intact and reminiscent of 14 nm thin layer at the periphery [2]. The aggregates for E161K showed a preferential clustering near the periphery, while in K97E the lamin aggregates were uniformly dispersed throughout the nucleoplasm. Line profiles of the lamin intensities across the nuclear cross-section showed two distinct peaks at the nuclear membrane in the WT, while interior peaks were observed in the two mutants (Fig. 1 C). Similar observations have been reported earlier in HeLa cells [19]. We performed western blots from the lysates of the cells and quantified similar levels of expression of endogenous lamin A/C as well as EGFP-lamin A proteins in ectopic manner (Fig. 1 D,E) thereby ruling out any possibilities of artefactual expression related effects. In the WT, the foci for the heterochromatin(HC) marker H3K9me3 were predominantly peripheral reflecting the known clustering of heterochromatic domains to the NL [20]. In contrast, we noticed a markedly different spatial organisation for both the lamin A and H3K9me3 foci in the mutants (Fig. 1 F). In E161K, the HC regions, marked in red in Fig. 1 F, were also perturbed from their predominant peripheral localisation, and could be found in the interior regions as well. This disruption was even more pronounced in K97E mutants, where a larger fraction of HC clusters were observed in the interior. To quantify these observations, we calculated the radial intensity profiles for both lamin A and heterochromatin for the WT and mutants (Fig. 1 G). As expected, in the WT, the lamin intensity shows a sharp peak at the periphery. The normalised intensity profile for HC represented a prominent peak just outside the peripheral lamin band, and subsequent smaller peaks could be observed in the interior as well. In E161K, although the peripheral lamin peak is still present, the relative intensity was much reduced compared to the WT, and smaller peaks are seen in the nuclear interior. The HC intensities again showed a larger signal outside the lamin peak. In K97E, the peripheral peak is completely disrupted resulting in a relatively uniform lamin signal throughout the nucleus. The HC peak at the nuclear periphery was also lost, with smaller peaks throughout the nucleoplasm. We next calculated the distribution of lamin proteins localized near the nuclear membrane and those present in the bulk nucleoplasm. In WT nuclei, a significant fraction of the lamin A was distributed uniformly in the nucleoplasm, consistent with earlier reports [20]. In contrast, this ratio of peripheral to bulk lamin reduced significantly by around 60% in the mutants E161K and K97E (Fig. 1 H) reinforcing the shift in organisation of lamin from the periphery towards the bulk. In the two mutants, we then characterised the ratio of lamin intensity signals inside the aggregates to free lamin in the nucleoplasm (Fig. 1 I). A much larger fraction of the nucleoplasmic lamin was found inside the aggregates in the E161K mutant. To investigate whether this is due to larger aggregate size or greater number of aggregates in E161K, we quantified the average radius of gyration (Fig. 1 J). The size of the lamin aggregates were relatively same in the two mutants, indicating that E161K had a larger number of aggregates compared to K97E. In addition, we measured the radius of gyration for the HC aggregates, demonstrating that the size of HC clusters decreased upon mutation, with similar HC-aggregate sizes in both mutants, albeit with a wider size distribution in E161K (Fig. 1 J).

**FIG. 1:**
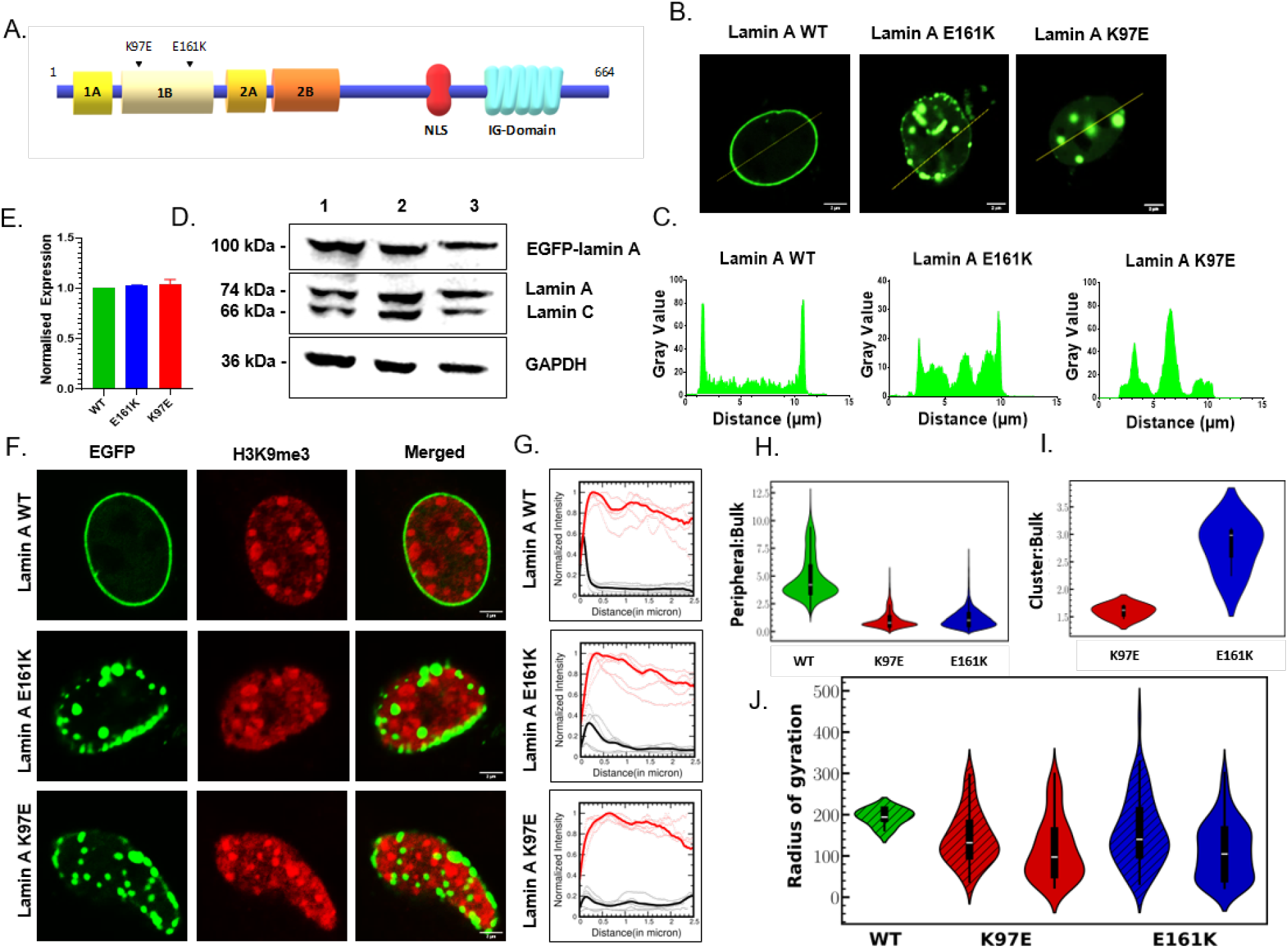
Lamin A mutants show disrupted lamin as well heterochromatin organisation. **A**. Schematic representation of lamin A, highlighting its rod domains 1A, 1B, 2A, and 2B, as well as the nuclear localization signal (NLS) situated between the rod domain and the Ig-domain. Black arrows indicate the positions of mutations K97E and E161K in 1B domain (Not to scale). B. Distribution of lamin A, visualized through confocal image of the nucleus (Scale bar 2µm). C. Intensity profiles across the nuclear axis depicting EGFP-lamin A distribution. D. Immunoblot against lamin A/C and GAPDH representing equal expression of EGFP-lamin A in C2C12 cells, 1) lamin A WT, 2) lamin A K97E, 3) lamin A E161K. E. The expression of EGFP-lamin A quantified from immunoblot done against lamin A/C, normalized to GAPDH. F. Confocal images depicting altered localization of heterochromatin marker H3K9me3. In the nucleus of cells expressing mutant lamin A, we can observe the shift of heterochromatin towards the center of the nucleus, as compared to peripheral localization of wild type (Scale bar 2µm). G. Normalised distribution of lamin A and heterochromatin marker H3K27me3, where lamin A is shown in Black curve and H3K27me3 is shown in Red curve. H. Violin plot representing distribution of lamin A as peripheral:bulk. I.Violin plot representing distribution of lamin A as cluster:bulk. J.Radius of gyration of lamin A and HC foci.

### B. Polymer models identify the key drivers behind lamin organisation

The experimental observations depicting large scale disruptions in both the lamin and chromatin organisation in the case of lamin A mutants suggest that chromatin organisation in WT nuclei is maintained through a sensitive balance between multiple interactions, including the lamin-lamin and lamin-heterochromatin interactions, as well as the lamin-membrane interactions. To understand the role of these interactions, we performed coarse grained polymer simulations for chromatin (Fig. 2 A). We modeled chromatin as a heteropolymer with EC (euchromatin) and HC (heterochromatin) beads (Fig. 2 B, see Methods for details).

**FIG. 2:**
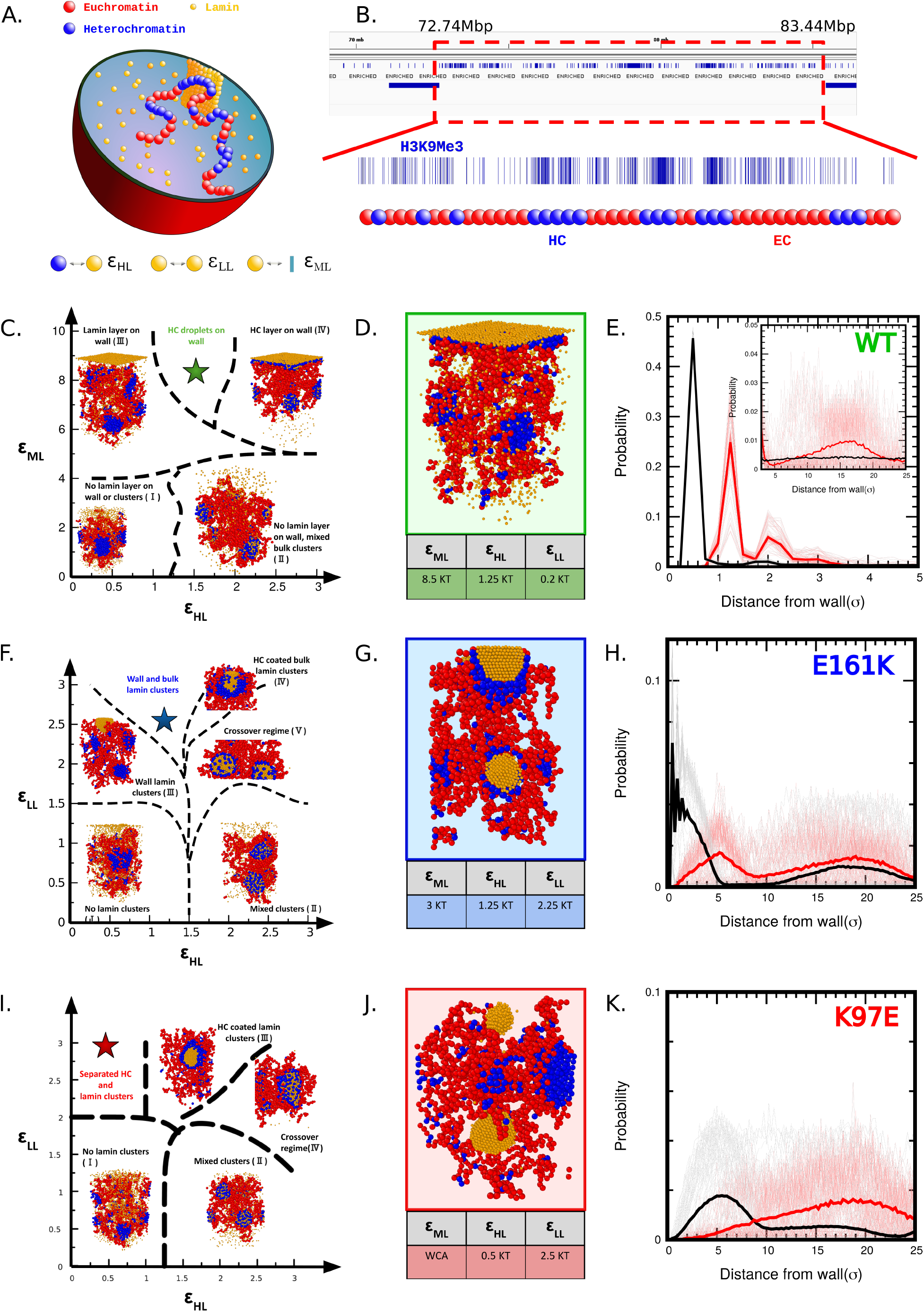
Polymer modeling of chromatin in presence of lamin. A. Schematic representation of our coarse-grained model illustrating chromatin-lamin interactions. Chromatin is modeled as a heteropolymer, where euchromatin (EC, red) and heterochromatin (HC, blue) beads. The lamin particles is depicted using yellow beads. B. Chromatin beads are classified as HC if the corresponding genomic region is enriched with the H3K9me3 modification, while all other regions are labeled as EC. C. A classification map of different structural configurations based on the interaction plane defined by the parameters *ϵ*_HL_ and *ϵ*_ML_ at low values of *ϵ*_LL_. D. A representative wild-type (WT) chromatin configuration from the region marked by a star in the interaction plane shown in panel C. E. XY-averaged z-density profiles showing the distribution of lamin (black) and HC (red) as a function of distance from the nuclear membrane. F. Classification map of configurations in the *ϵ*_HL_ *− ϵ*_LL_ interaction plane for a fixed *ϵ*_ML_ value of 3.0. G. A representative configuration of the E161K mutant from the region marked by a star in the interaction plane shown in panel F. H. XY-averaged z-density profiles of lamin (black) and HC (red) for the E161K mutant, plotted as a function of distance from the nuclear membrane. I. Configuration classifications for different states in the *ϵ*_HL_ *− ϵ*_LL_ interaction plane when *ϵ*_ML_ is set to the purely repulsive Weeks-Chandler-Andersen (WCA) potential. J. A representative configuration of the K97E mutant, corresponding to the star-marked region in the interaction plane shown in panel I. K. XY-averaged z-density profiles of lamin (black) and HC (red) for the K97E mutant, showing the distribution relative to the nuclear membrane.

Since we are also interested in the lamin organisation, we adopted a coarse grained particle representation for lamin proteins as well (Fig. 2 A). This realistic representation of lamins as discrete particles is a departure from previous models that treated the nuclear lamina as a uniform surface layer, providing a more detailed and accurate description of lamin-chromatin interactions in our simulation. The interactions between EC and HC, EC and EC, EC and lamin, and between HC and HC regions were assumed to be fixed throughout this manuscript, consistent with parameter estimated from earlier studies (see Methods for details) [56, 57, 59].

We performed the simulations within a confined geometry to study the emergent organisation depending on the underlying interaction parameters. We focused on the role of three interactions - the preferential interaction between heterochromatin and lamin (ϵ_HL_), the interaction between lamin beads (ϵ_LL_) and between the nuclear membrane and lamin (ϵ_ML_).

#### 1. Weak lamin-lamin interaction

For WT nuclei, lamin proteins were uniformly distributed underneath the nuclear membrane thereby indicating the presence of a strong lamin-membrane attractive interaction (ϵ_ML_) (Fig. 1 B). In contrast, there were no significant lamin aggregates in the bulk of the nucleus. We chose *ϵ*_LL_ = 0.2 (=*ϵ*_EE_) to model these weaker lamin-lamin interactions in the wild type. HC regions were found to be clustered throughout the nucleus, with a higher probability of clustering near the nuclear membrane due to interactions between lamin and regions marked by H3K9Me3 modifications (ϵ_HL_)(Fig. 1 F,G,H). For these WT nuclei then, we explored the parameter space encoded by these two energy scales - *ϵ*_ML_ and *ϵ*_HL_ - to determine where the WT phenotypes lie in this interaction plane. The classification of possible configurations is shown in Fig. 2 C. For low values of *ϵ*_ML_, the attraction to the membrane was not sufficient to form a lamin layer at the periphery. If the interaction between HC and lamin was also low, lamins remained dispersed throughout the bulk (configuration I). As *ϵ*_HL_ increased for low *ϵ*_ML_, lamins started forming aggregates in the bulk due to effective attractive interactions mediated by heterochromatin (configuration II). We denoted the lamin aggregates in this region as mixed aggregates, since the any single aggregate contains large fractions of both lamins and HC. There were no distinct HC clusters in this regime. Now, as *ϵ*_ML_ increased, lamins started migrating to the periphery. Beyond a threshold *ϵ*_ML_, we observed the formation of lamin layer at the boundary. This threshold was slightly higher for larger HC-lamin interaction strengths since lamins would now have to overcome the attraction to the HC regions to migrate to the periphery. For high *ϵ*_ML_ and low *ϵ*_HL_, we observe a lamin surface layer with HC clusters dispersed throughout the bulk (configuration III). For all these three configurations, we observed no preferential clustering of HC domain near the periphery. On the other hand, for very large *ϵ*_HL_, the HC spread throughout the surface through a wetting-like transition, and HC clusters in the bulk also possessed significant lamin fractions (configuration IV). For intermediate values of *ϵ*_HL_, we observed a lamin layer at the surface, HC clusters at the surface and the bulk, and no mixed lamin and HC aggregates, all of which corroborated to the experimental WT observations (Fig. 1 F,G). We thus identified this region of parameter space as corresponding to the WT, denoted by a star in Fig. 2C. In our coarse-grained model, we thus chose *ϵ*_ML_ = 8.5 and *ϵ*_HL_ = 1.25 to model our WT nuclei. This implied the hierarchy of energy scales *ϵ*_EE_ < *ϵ*_HL_ < *ϵ*_HH_ [56, 57]. The corresponding representative snapshot is shown in Fig. 2 D. The radial distribution profiles for both lamin and HC are plotted in Fig. 2 E, recapitulating the peripheral lamin organisation, and a preferential clustering of HC near the membrane in addition to dispersed HC clusters in the interior.

### 2. Weak lamin-membrane interaction

The E161K mutant introduced distinct perturbations in the chromatin organization, with marked changes in the distribution of lamin proteins. The peripheral lamin organisation seen in WT was largely disrupted, and there was a simultaneous increase in lamin aggregate formation, indicating a lamin-lamin interaction (*ϵ*_LL_). The lamin aggregates showed a tendency to cluster near the periphery, although they were also present throughout the nucleoplasm, indicating a reduced (compared to WT) but finite *ϵ*_ML_ interaction. We further observed a preferential localisation of the HC clusters towards the periphery, suggesting a significant HC-lamin interaction (*ϵ*_HL_). Based on our analysis in Fig. 2 C, we chose *ϵ*_ML_ = 3, such that this interaction lies below the threshold for a surface lamin layer for any value of *ϵ*_HL_. We now proceeded to explore the possible configurations in the *ϵ*_HL_-*ϵ*_LL_ plane for *ϵ*_ML_ = 3 (Fig. 2 F). For low *ϵ*_LL_ and low *ϵ*_HL_, we observed no lamin aggregates (configuration I). On increasing *ϵ*_HL_, we observed the formation of HC mediated mixed lamin aggregates (configuration II). Next, on increasing *ϵ*_LL_ for low values of *ϵ*_HL_, we observed distinct lamin and HC clusters (configuration III). The lamin aggregates in this case were preferentially located near the periphery due to the *ϵ*_ML_ interaction. On increasing the *ϵ*_HL_ strength and for relatively high values of *ϵ*_LL_, lamin aggregates continued to be formed near the periphery, but there were also lamin aggregates in the bulk due to direct interactions. Further, the relatively higher *ϵ*_HL_ resulted in increased co-localisation of the HC clusters near these lamin aggregates. This region of the interaction plane then corresponded to our E161K phenotype and is marked with a star in Fig. 2 F. For even higher strengths of *ϵ*_HL_, we observed a new configuration - HC coated lamin aggregates (configuration IV). In this regime, the lamin-lamin interactions were strong enough to form aggregates via direct interaction, and the *ϵ*_HL_ interaction was not sufficient to dislodge the lamins from the cluster interior, thus forming a coating layer of HC outside the lamin aggregate. On increasing the *ϵ*_HL_, or decreasing the *ϵ*_LL_, these two energy scales competed, driving the penetration of a fraction of HC into the lamin aggregates, resulting in a crossover regime (configuration V). For the E161K phenotype then, we chose *ϵ*_ML_ = 3, *ϵ*_HL_ = 1.25, and *ϵ*_LL_ = 2.25. The corresponding configuration is shown in Fig. 2 G. The corresponding lamin and HC probability distributions are shown in Fig. 2 H which qualitatively recapitulated the radial distributions seen experimentally in Fig. 1 G.

### 3. Disrupted lamin-membrane interaction

The K97E mutation again significantly affected both chromatin and lamin organization. The peripheral lamin organisation along the nuclear membrane was largely depleted, which we modelled through a repulsive WCA lamin-membrane interaction (*ϵ*_ML_ = WCA). Simultaneously, lamin aggregates emerged in the bulk (Fig. 1 B), indicating a higher degree of lamin-lamin interactions compared to the WT. Further, in contrast to E161K, there appeared to be no preferential colocalisation between lamin and HC clusters, indicating a disrupted lamin-HC interaction (*ϵ*_HL_). For the K97E phenotype then, we explored the space of possible configurations encoded by the two interaction energies - *ϵ*_LL_ and *ϵ*_HL_. This is shown in Fig. 2 I. For low values of *ϵ*_LL_ and *ϵ*_HL_, no lamin aggregates formed (configuration I). However, on increasing the HC-lamin interaction, lamins could aggregate via effective attractive interactions mediated by HC, as in the WT case, resulting in mixed aggregates (configuration II). For high *ϵ*_LL_ and low *ϵ*_HL_, we again observed the formation of lamin clusters. These were pure lamin aggregates formed by direct interaction. However, this threshold *ϵ*_LL_ was higher compared to Fig. 2 F, since in this case there was no *ϵ*_ML_ interaction to assist lamin aggregation. In this regime of interaction space denoted by a star in Fig. 2 I, lamin and HC clusters were distinct, and thus represented the phenotypes seen experimentally for K97E. For these high values of *ϵ*_LL_, on increasing the HC-lamin interaction strength, we again observed the HC coated lamin aggregates (configuration III), which transitioned to the crossover regime (configuration IV) on decreasing *ϵ*_LL_ or increasing *ϵ*_HL_. For the K97E phenotype then, we choose *ϵ*_LL_ = 2.5 and *ϵ*_HL_ = 0.5, which resulted in distinct lamin and HC clusters dispersed throughout the system (Fig. 2 J). The probability plots for the lamin and the HC distributions are shown in Fig. 2 K.

In summary, our polymer models suggest that lamin-mediated chromatin organisation was maintained through an interplay of the lamin-lamin, lamin-membrane, as well as lamin-heterochromatin interactions. Perturbing these interactions through mutations could then lead to widespread disruption of spatial organisation.

### C. Lamin A mutants K97E and E161K show differential 3D repositioning of heterochromatin and altered lamin-chromatin colocalisation

Our polymer models suggested that lamin-mediated chromatin organization is maintained through an interplay of the lamin-lamin, lamin-membrane, as well as lamin-HC interactions. It also indicated that perturbation of these interactions in K97E and E161K mutations might be the cause of widespread disruption of spatial HC organization. To verify the simulation results of the strong lamin-lamin interactions in the mutants coupled with weak lamin-chromatin interaction, we performed 3D fluorescence in-situ hybridization (3D-FISH) experiment to examine the localization of chromosome 13 (Chr. 13), which has a heterochromatic short arm [82] and localised essentially towards nuclear periphery in cells expressing wild-type lamin A, as was shown in the top row of Fig. 3 A. For the mutants, as shown in the middle and bottom rows this peripheral organisation of chromosome 13 was lost, simultaneously with the loss of the peripheral lamin layer and the formation of lamin aggregates. We first investigated the the interaction of chromosome 13 with the EGFP-lamin A, by a colocalisation analysis from the images (right two panels of Fig. 3 A). The overlap between the lamin and chromosome 13 signals was strongest in the WT cells, and was significantly reduced in E161K and almost non-existent in K97E mutants. We quantified this by calculating Pearson’s correlation coefficient between chromosome 13 and EGFP-lamin A signals (Fig. 3 B). The values of Pearson’s Coefficient for WT, E161K and K97E are 0.26, 0.09 and 0.04 respectively, which pointed to a decreased lamin-chromatin colocalisation events in mutants in comparison to wild type lamin A expressing cells, the least co-localization being in K97E (Fig. 3 B). This data reported for the first time a decreased lamin-chromatin co-localization in mutants in comparison to wild type lamin A expressing cells.

**FIG. 3:**
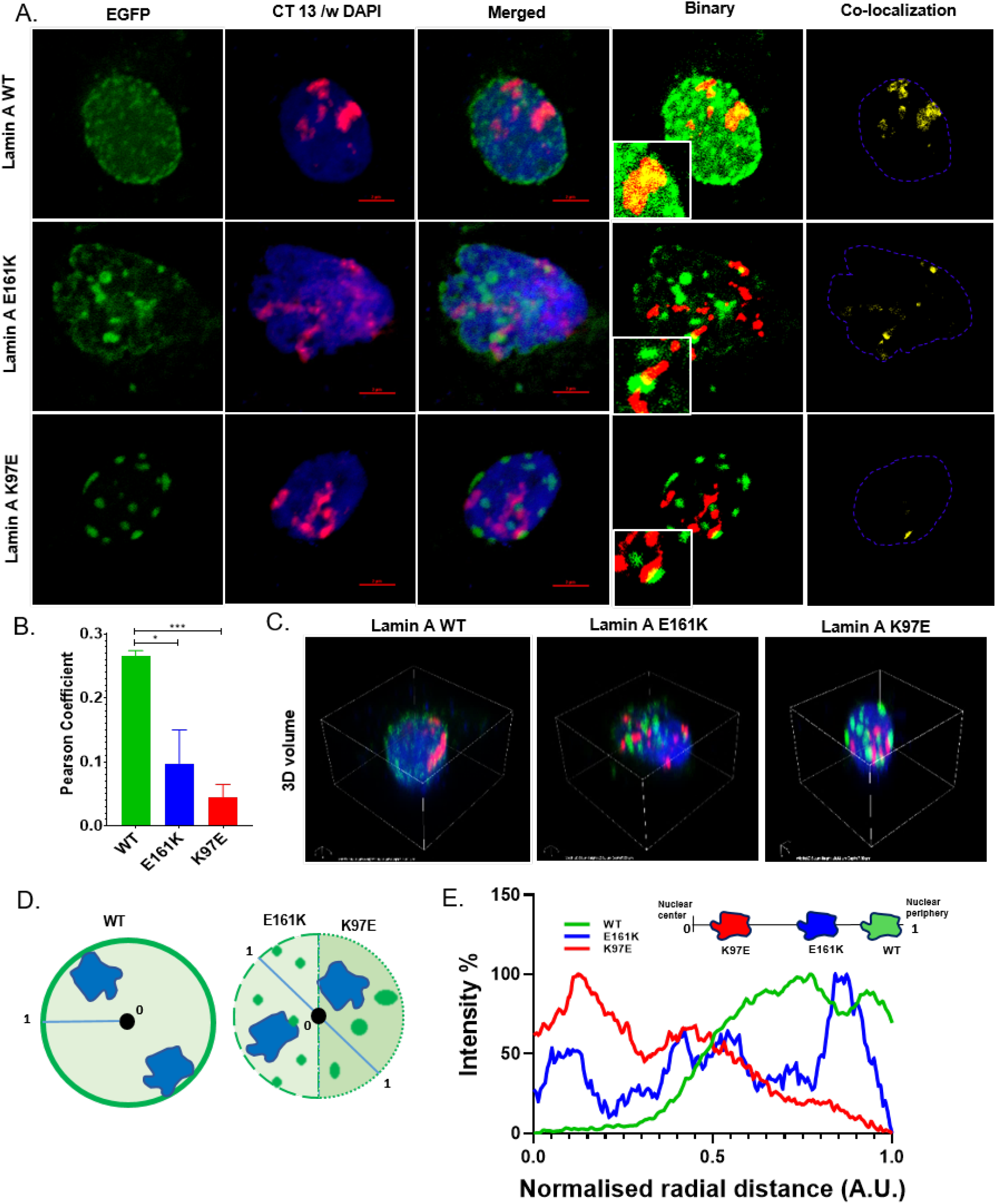
E161K and K97E mutation in lamin A results in a territorial shift of Chr. 13 and induces a reduced lamin-chromatin interaction. A. Representative FISH experiments showing distribution of Chr. 13, in nucleus of Hela cells transfected with pEGFP-lamin A WT and mutants. Maximum intensity projection of confocal image stack with three color channels, illustrates the distribution of lamin A (green), along with position of Chr. 13 territory (red) in a nuclear volume stained with DAPI (blue). Composite binary images showcase the regions of overlap between lamin and chromosome territory (yellow). The last row marks the regions of colocalization within the nucleus from the composite image (Scale bar 2µm). B. Pearson’s Coefficient between lamin A and Chr. 13. C. Representative 3D volume snap depicting Chr. 13 territory within the nuclear volume. D. Schematic of the model adopted for 3D distance measurements of Chr. 13 in blue, with normalised radial distance from geometric center of the nucleus (0) to the nuclear periphery (1), in three different scenarios of lamin expression. E. Normalised radial distance of Chr. 13 is plotted for WT, K97E and E161K lamin A transfected cells.

The 3D-FISH experiments pointed to large-scale alterations in the positioning of chromosome 13 in the mutants. This can also be seen from the representative 3D-volume rendition images from the wild type and mutant transfected cells depicted in Fig. 3 C for chromosome 13 and lamin A proteins in 3D nuclear space. To quantify the effect of mutant lamin A proteins on positioning of chromosomes, we checked for the radial distribution of chromosome 13 in cells exhibiting ectopic expression of WT, E161K and K97E proteins. Fig. 3 D depicted the model used for calculating the radial distance of the Chr. 13 territories. We found a distinct shift in the distribution of Chr. 13 from the periphery towards the centre in both the mutant proteins, most pronounced being that of K97E (Fig. 3 E). This was consistent with the results of the colocalisation analysis, where, the maximum disruption of lamin-chromosome 13 colocalisation was observed for the K97E mutant. This altered positioning of chromosome 13 was also in agreement with previous results in cardiomyopathic E161K mutation, where chromosome 13 migrated towards nuclear interior from its designated peripheral position, whereas in other mutations such as *LMNA* Δ303 or D596N which were derived from patients with inherited familial cardiomyopathy chromosome 13 showed a further peripheral relocation towards the extreme edge of nucleus [83]. Interestingly, differential alterations of 3D radial distribution of chromosome territories were reported in various pathological conditions caused by different *LMNA* mutations [84–88]. Also, the fact that altered chromosomal positioning is absent in laminopathy patient lymphoblasts, a cell-type which shows minimal expression of A-type lamins [89] is another indication of involvement of lamin in determination of chromosome territories.

To summarise, we reported for the first time a decreased lamin-chromatin colocalization in mutants and deciphered quantitatively that the radial distribution of chromosome 13 changed in the K97E and E161K in comparison to the WT, consistent with a significant decrease in lamin-chromatin interaction. As the lamin-lamin interactions increased in the mutants, giving rise to lamin aggregates, there is a simultaneous increased segregation between the lamin and the HC clusters.

### D. Lamin A mutants K97E and E161K form liquid liquid phase separation inside nucleus

Our analysis so far indicates that the structural changes in the mutated lamin proteins leads to decreased lamin-chromatin interaction in favour of more pronounced lamin-lamin interactions, leading to aggregate formation. These changes in structure leading to global changes in conformations should also affect the dynamics of lamin inside the nucleus. In order to investigate this, we examined the distribution and dynamics of lamin A in the nuclear milieu by following the spatio-temporal envelope of lamin A in the nucleus of the cell through a series of FRAP experiments. Previously, various groups have elucidated the dynamic properties of lamin A inside the nucleus by FRAP experiments [90, 91]. But none of these experiments could show the spatio-temporal dynamics of mutant lamin A which form aggregates, and thus could not distinguish between the dynamics of nucleoplasm and aggregates independently. In contrast, we designed our experiments in a manner that allowed us to clearly distinguish between the dynamics of nucleoplasmic lamin and the lamin present in the interior of the aggregates of lamin A mutants. The aggregates in the mutants have typical areas of more than 2.39 *µ*m^2^ [92], and hence we chose a bleach area to 0.25 *µ*m^2^ to confine the ROI strictly inside the aggregate, in order to distinguish the recovery events from the aggregate itself or the nucleoplasm. We selected cells expressing equal and moderate amount of the EGFP-lamin A proteins for our study (Fig. 1 B). The photo-bleached regions of lamin A WT and the mutants K97E and E161K were analysed for recovery as a function of time, for three distinct nuclear regions – (i) in the bulk nucleoplasm, (ii) inside the lamina(WT) or aggregates(K97E/E161K), and a (iii) mixed region corresponding to a 50:50 distribution of inside and outside the aggregates (Fig. 4 A-C). The bleaching and recovery curves for these three scenarios are shown in Fig. 4 D-F. The mobile fractions of the lamina/aggregate, nucleoplasm and mixed regions has been shown in Fig. 4 G. The WT rim or the lamina corresponded to the least mobile fraction of 3.98% whereas the aggregates for the mutants have much larger mobile fractions of 22.37% and 17% for K97E and E161K respectively, suggesting a fluidic nature of the aggregates. As expected, the nucleoplasmic pools consisted of large mobile fractions corresponding to 36.02% for WT and 60.4% and 44.8% for K97E and E161K respectively (Table II). 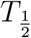 values showed that wild type lamin A had two principal distribution profiles – one as a distinctly immobile phase in the nuclear rim with a 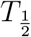 of 43.9 sec and the other a mobile phase in the nucleoplasm with 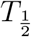 of 7.3 sec (Fig. 4 H, Table II). For nucleoplasmic lamin A, the slowest recovering state belonged to the wild type protein while K97E recovered the fastest that was reflected in the 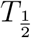 value of 4.21 sec for K97E and 4.81 sec for E161K mutant (Fig. 4 H, Table II). Focusing on the dynamics of mutant aggregates, we found that both the mutants show an internal rearrangement, with a 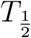 of 27.36 and 32.42 sec for K97E and E161K respectively (Table II). These observations were also borne out by the comparison of estimated diffusion constants of WT with the mutants, where we observed a faster diffusion rate for the mutants in all three regions, with K97E being the fastest diffusing species(Fig. 4 I). The lower mobile fractions and slower recovery times in the lamin aggregates in E161K as compared to K97E can be understood through the polymer simulations and the colocalisation analysis. In E161K, HC domains are preferentially located close to the lamin aggregates, and this association with the polymer can slow down the dynamics of lamin as well as lead to smaller mobile fractions. Although these findings seemed to be counter-intuitive at first glance in terms of the aggregation events [19], we envisaged it in the light of lamin-chromatin interaction that might be abrogated in the mutants especially K97E in favour of elevated lamin-lamin interaction. This is corroborated by the separation of the lamin A and HC foci. In summary, our FRAP experiments demonstrate that there is both internal rearrangements within the aggregates, as well as exchange between the aggregates and nucleoplasmic pools of lamin A, emphasizing the fluidic nature of the aggregates.

**FIG. 4:**
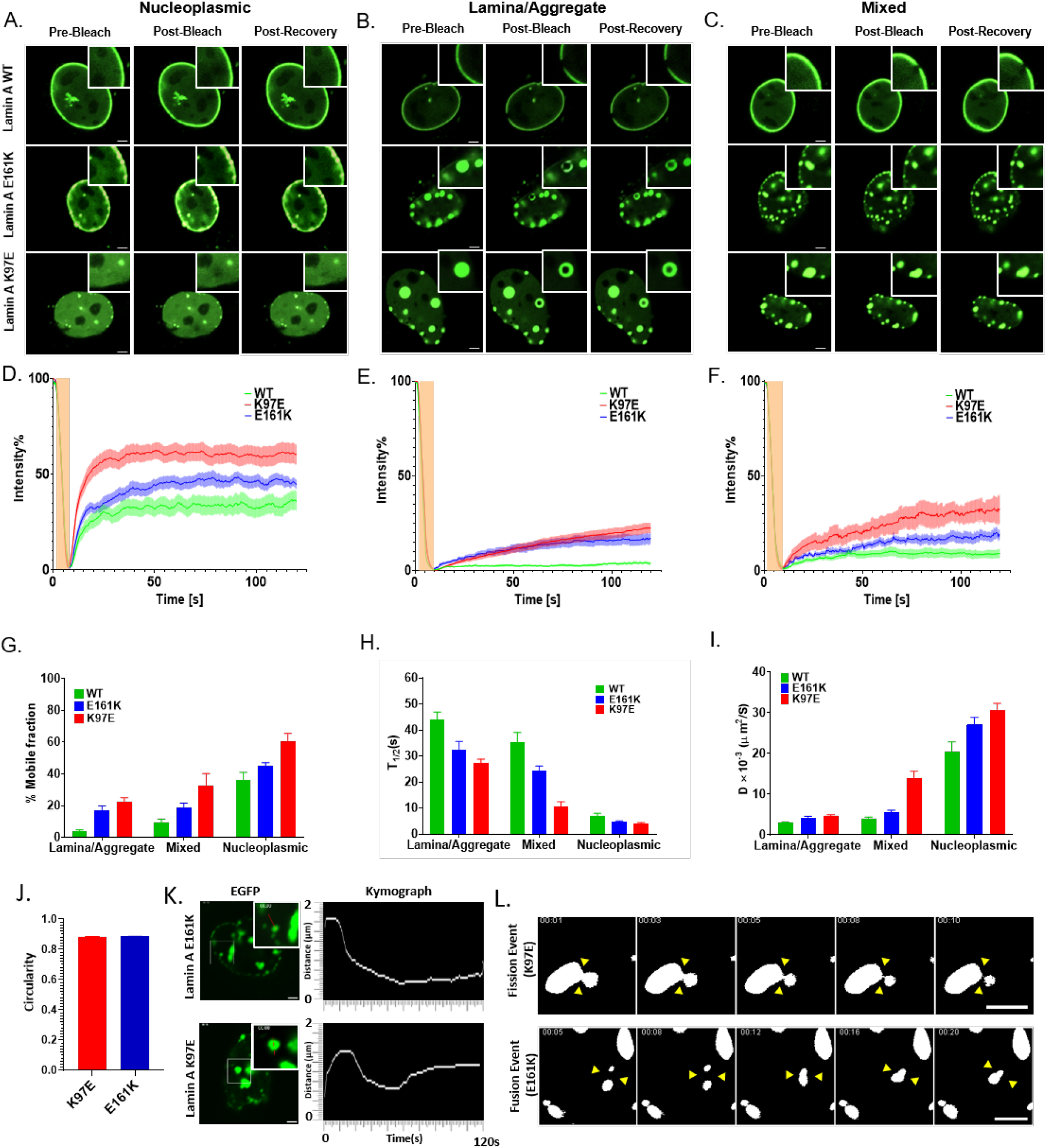
Dynamics of lamin A diffusion within nuclear aggregates exhibit behavior reminiscent of liquid-liquid phase separation (LLPS). A, B, and C. Fluorescent Recovery After Photobleaching (FRAP) confocal images captured pre-bleach, post-bleach, and post-110-second recovery period for EGFP-lamin A. A. Bleaching events within a purely nucleoplasmic ROI. B. Bleaching events with the lamina/aggregates. C. A balanced ROI consisting of mixed region within and outside the aggregates, (approximately 50-50), underscoring the mixed dynamics of lamin A in both wild-type and mutant. All insets provide zoomed-in views of the ROIs (Scale bar 1*µ*m). D, E, and F. Depiction of the recovery of fluorescent intensities within the aforementioned ROIs during the bleaching experiment, as a function of time, with bleaching event marked with yellow. G. The mobile fraction of lamin A within ROIs of wild-type and mutant cells across the three regions. H. The calculated half-time 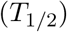 in both wild-type and mutant contexts. I. The diffusion constants calculated with respect to the half-life of fluorescent recovery. J. A representation of the average circularity of lamin A mutant condensates, illustrating their nearly circular appearance. A perfect circle is indicated by a circularity value of 1. K. A skeletonized kymograph showing the movements of mutant condensates within the nucleus over a 120-second time frame. L. Representative montage of binarized images, generated from time-lapse imaging of EGFP-lamin A mutants, showing fission and fusion events of mutant condensates (represented by yellow triangles). Error bars represent SEM in all the graphs.

This property of internal rearrangement and external exchange is in fact one of the hallmarks of liquid-liquid phase separation (LLPS) [93, 94], suggesting that formation of lamin aggregates in the mutants proceeds through LLPS. In order to verify this, we focused on the other key properties that define LLPS - the spherical nature of the aggregates, and the fusion of smaller aggregates to form larger ones [93]. We found out that the average circularity (2D equivalent of sphericity) of both K97E and E161K mutants were close to 0.9, where the circularity of a perfect circle is 1 (Fig. 4 J). The spherical nature of smallest surface area is one the most common physical attributes of fluidity and our characterization of the lamin A mutant aggregates confirm this liquid-like property. Observations of fusion to form larges condensates in biological LLPS is itself a very rare occasion due to the heterogeneous nature and long timescales involved. A primary requirement for the fusion of two droplets or condensates is that they need to come into proximity and hence should have a detectable mobility on a cellular timescale. To check if the condensates were mobile, we performed a time lapse imaging of the mutant condensates for 120 sec and plotted as a kymograph of the moving condensates (Fig. 4 K). From the kymographs we could observe that both K97E and E161K condensates exhibited a mobility of approximately 1-1.5 *µ*m in less than a minute. This mobility increased the probability of contact hence that of fusion of the condensates. A representative fusion event where two droplets meet and merge into a single larger droplet is shown in Fig 4 L. Interestingly, during the time lapse imaging, in addition to fusion events, we also captured fission of large droplets into two smaller ones (Fig 4 L). Fission of droplets is unexpected in the context of equilibrium phase separation theories, and suggests that non-equilibrium forces may play a role in the kinetics of these lamin A mutant condensates. Through the observations regarding internal dynamics of lamin A mutant condensates, their spherical nature and mobility, we believe that this is the first report showing condensate formation of mutant lamin A proteins akin to LLPS between the nucleoplasm and condensates.

## III. DISCUSSION

The interaction between lamin A and chromatin is crucial for chromatin positioning as well as epigenetic regulation [2, 55, 95]. The peripheral distribution of lamin A in WT cells regulates the spatial organization of chromatin. Our study highlighted the functional aspects of lamin A, underscoring its interactions with both the nuclear membrane and HC. We found that K97E and E161K mutants of lamin A protein significantly disrupted these interactions, leading to an aberrant chromatin positioning and lamin aggregation. In both mutants, we observed drastic disruption in the peripheral organisation, together with the formation of large nucleoplasmic condensates, indicating enhanced lamin-lamin interactions coupled with weakened lamin-chromatin association. We performed polymer simulations to understand these observations. Crucially, in a departure from previous models of lamin mediated chromatin organization, we adopted a coarse-grained particle model for lamins, allowing us to investigate the effect of nucleoplasmic as well as membrane-bound lamin on the spatial organisation. Our polymer simulations demonstrated the delicate balance of lamin-lamin, lamin-chromatin, and lamin-membrane interactions that is essential for maintaining proper HC organization in wild-type cells. We obtained a broad classification of configurations in the plane of these different interactions, and the different phenotypes observed in the simulations might serve as a template to understand aberrant organisation in lamin mutants beyond the K97E and E161K mutants studied here.

HC regions of DNA have been shown to be organized inside the nucleus as a phase separated compartment with the help of intrinsically disordered region (IDR) of HP1 proteins, creating a partial LLPS inside nucleus [93, 94, 96, 97]. Similarly, lamins also have a long stretch of disordered region in its tail domain except the Ig-fold domain [98, 99]. As a sequel to the mutations, the helicity of the rod domain decreases [19, 92] thereby augmenting the effect of the disordered region in the process of the assembly. This increased disordered stretch with altered secondary and tertiary structures of the proteins might be the contributing factor for the LLPS like behaviour of mutants. It must be emphasized that LLPS in cellular systems result from weak, multivalent, and non-covalent interactions between proteins and nucleic acids [100]. These low affinity interactions tend to increase the self-associations between the macromolecules thereby lowering the entropy compared to macromolecules-solvent interactions. Interestingly, our finding on LLPS in the mutant condensates is strongly supported by the thermodynamic data obtained from ITC experiments where the change in entropy Δ*S*^WT^>Δ*S*^E161K^>>Δ*S*^K97E^ [19]. Focusing on the lamin aggregates in the mutants, for the first time we showed qualitatively that mutant lamin A proteins exhibit altered dynamics within the nucleus, with characteristics reminiscent of LLPS. The spherical nature of the aggregates along with their internal rearrangements and fusion of smaller droplets confirmed a liquid-like phase. In addition, fission events suggest a possible role for out-of-equilibrium dynamics in determining the kinetics in these systems. Our cumulative findings from confocal imaging, coarse-grained simulations, 3D-FISH and FRAP experiments indicated that lamin A mutants underwent LLPS within the nucleus potentially due to weakened, multivalent interactions, aligning with the behaviour observed in other nuclear structures exhibiting LLPS. These disruptions are indicative of potential implications for diseases such as DCM, where chromatin organization as well as epigenetic regulation is severely compromised. These findings open an avenue for exploring the broader impact of these mutations with a new and versatile perspective which is able to explain chromatin repositioning during the diseased conditions.

## IV. MATERIALS AND METHODS

### A. Site-Directed Mutagenesis

Site-directed mutagenesis was carried out in pEGFP-lamin A following standard protocol as detailed in previous articles from our lab [19]. The wild type lamin A and the mutants are denoted byLamin A WT or WT, lamin A K97E or K97E, and lamin A E161K or E161K respectively in all subsequent experiments.

### B. Cell Culture and Transfection

C2C12 (ATCC) and HeLa (ATCC) cells were maintained in ATCC formulated DMEM supplemented with Penicillin-streptomycin as mentioned earlier [19, 64, 92]. Cells were cultured using a standard protocol and maintained at 37°C and 5% CO_2_. Cells between 2nd and 3rd passages were seeded to a confluency of 50 *−* 60% for transfection. 3*µ*g of DNA was used for each transfection using Lipofectamine 3000 (Invitrogen, USA) according to manufacturers protocol. Cells were processed for further experiments after 24 hours of transfection.

### C. Western Blotting

Protein concentration was estimated by Bradford assay as described earlier [19, 101, 102]. Equal volumes of protein samples and Laemmli buffer were mixed and boiled at 100°C for 10 min prior to gel loading. Proteins were resolved using 10% SDS gel electrophoresis and transferred onto a nitrocellulose membrane (Merck Millipore, USA) and the subsequent steps of the western blot were performed according to the methods reported earlier [18]. Primary antibodies used in this study were Anti- lamin A/C Monoclonal Antibody (L2193, Thermo Fisher, USA), and Anti-GAPDH antibody (ab9485, Abcam, USA) at 1:500 and 1:1000 dilutions respectively in PBST/milk or PBST. HRP conjugated Goat-Anti Rabbit (32460, Thermo Scientific, USA) was used as the secondary antibody. The blots were thenexposed to enhanced chemiluminescent substrate (Thermo Scientific, Pierce, USA) and imaged on a chemiluminescence detector (Azure Biosystems 280, USA). Subsequently band intensities were quantified using Image J software (ImageJ bundled with 64-bit Java 1.8.0.112).

### D. Immunofluorescence staining

After 24 hours of transfection, cells were fixed with 4% paraformaldehyde for 15 minutes. Fixed cells were permeabilized with 0.5% Triton X-100 for 5 minutes, and incubated with primary antibody solution containing blocking agent (5% Normal Goat Serum in PBS) for 2 hours at room temperature. Following incubation with primary antibody, cells were then washed thrice with 0.05% Tween-20 in PBS followed by thrice with PBS at room temperature, after which the cells were treated with secondary antibody diluted in PBS for 2 hours at room temperature. Cells were mounted with Vectashield Plus (Fisher Scientific, USA) containing DAPI. Primary antibody dilution for anti H3K9me3 antibody (D4W1U, Cell Signalling Technology, USA) was 1:100. Secondary antibodies were conjugated with Alexa Fluor 546 with a dilution of 1:400.

### E. Confocal imaging

For confocal imaging, the slides were visualized by 100X oil DIC N2 objective 1.40NA/1.515 RI in NIKON TiE inverted research microscope with a digital 4X zoom. The images were captured in resonant mode. The excitation filters used were 450/50, 525/50,595/50, and the first dichroic mirror used was 405/488/561. The lasers used were Multi-line Argon-Krypton mixed gas laser λ 488nm, Solid state laser λ 405nm and, Solid state laser λ 561nm. For 3D-FISH imaging the resonant scanner with a scanner zoom: 6.0 and line averaging: 4.0 was used. While capturing the images, pinhole was maintained at 69.0 m, one way scan direction and channel series mode line 1-4 was used. The Z-stack imaging was carried out in a step size of 0.2 *µ*m.

### F. Image analysis

Images were processed using Nis Elements Analysis AR (Ver 4.13) and ImageJ software (Ver 1.8.0 112). To gain insights into lamin and heterochromatin organisations, we computed radial intensity profiles for both lamin and heterochromatin. We determined the center of the nucleus and gathered linear profiles of lamin and HC along the line joining a point on periphery to center. We then averaged these profiles along all possible direction to obtain theta averaged intensity profiles. In addition to overall lamin distribution, we investigated the organization of lamins by assessing the ratio of clustered to non-clustered lamin proteins in samples. This ratio revealed whether lamins tended to aggregate in specific regions or were more evenly dispersed within the nucleus. We also examined the relative positions of heterochromatin clusters and lamin clusters within the nucleus. By measuring the distances of these clusters from the nuclear boundary, we were able to determine how these important structures were positioned in relation to the nucleus’s periphery. By conducting this multifaceted analysis, we aimed to provide a comprehensive understanding of the differences between WT and mutant nuclei in terms of lamin distribution, clustering behavior, and the spatial organization of heterochromatin and lamin clusters within the nucleus. This data contributes to a deeper exploration of nuclear structure and function in the context of genetic mutations.

Pearson’s Coefficient and Overlap coefficient were calculated from the using JACoP plugin [103] in ImageJ utilizing the following formulae:

Pearson’s coefficient:

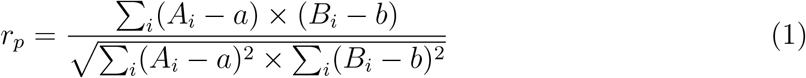

Where, Ai and Bi corresponds to grey values of i-numbered voxels in channel A and channel B respectively; the corresponding average intensities over the full image is noted as a and b.

The circularity of the condensates was measured using the ‘Analyze Particles’ function in ImageJ. This tool provided an assessment of the shape of the condensates, allowing the quantification of their circularity based on their projected area and perimeter with the help of the formula:

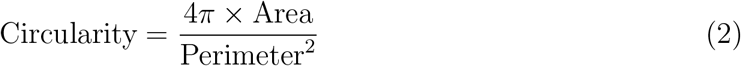

Additionally, a kymograph was generated using the ‘Multiple Kymograph’ function in ImageJ, with a ROI set at 2 *µ*m, with line width of 1 pixel. In the resulting kymograph, the y-axis represented the spatial position of the line ROI, while the x-axis represented the time progression. This visualization enabled tracking of the dynamic behaviour of the condensates over time. To further analyze the movement and structural features of the particles, the kymograph was processed using the ‘Analyze Skeleton’ function. This allowed for the extraction of the trajectory of the particles within the kymograph, providing detailed information on their motion throughout the time-lapse experiment.

### G. Details on the setup and simulations of the coarse-grained model

#### Coarse-grained chromatin and lamin

We constructed a model of a specific region on Mouse chromosome 18 (GRCm38/mm10), covering 10.7 Mbp. In this coarse-grained approach, the chromatin is represented as a flexible bead-spring heteropolymer chain made up of 2140 beads, each with a diameter of *σ*. Each bead corresponds to a 5-kilobase pair (kbp) segment of DNA, approximately equivalent to 25 nucleosomes. The beads were classified into two types: euchromatin (EC), shown in red, and heterochromatin (HC), shown in blue. This classification was based on chromatin immunoprecipitation sequencing (ChIP-seq) data for the H3K9me3 mark. Beads were labeled as HC if their corresponding genomic region contained a peak in the H3K9me3 signal.

#### Simulation box

To accurately capture the realistic chromatin density within the nucleus, we configured the simulation to run inside a rectangular box with dimensions of 20*σ* in the x and y directions and 35*σ* in the z direction. Periodic boundary conditions were applied in the x and y directions, effectively creating an infinite grid of repeating boxes. In the z direction, however, we used fixed boundaries to simulate confinement, reflecting a small region of the nucleus adjacent to the nuclear membrane. We considered a total of 2140 chromatin beads in our simulation. Now, the volume fraction of the chromatin:

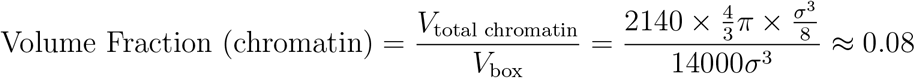

We also randomly placed 2000 coarse-grained lamin beads in the box, where each lamin bead has a diameter of *σ*/2. Thus, the lamin volume fraction is then:

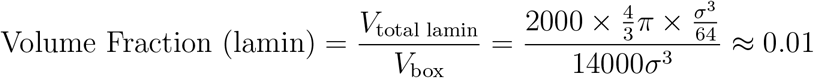

#### Interactions between chromatin and lamin beads

The total energy (U) of the bead-spring polymer is expressed as:

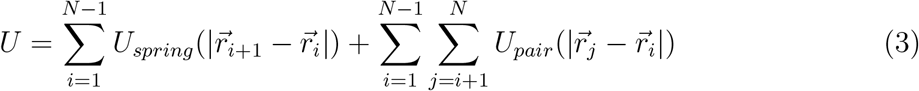

Where 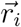 denotes the position vector of the *i*^*th*^ bead. The first term describes the polymer’s connectivity via harmonic springs, given by:

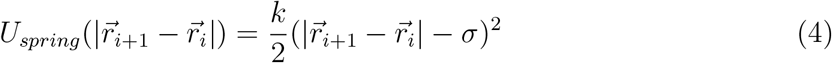

Here, *k* represents the spring constant, with a value of *k* = 60*k*_*B*_*T*. In addition to this, short-range interactions—both attractive and repulsive—are included for beads that come close to each other in 3D space. The second term in the total energy equation accounts for these non-bonded pairwise interactions, modeled through either the Lennard-Jones (LJ) potential or a purely repulsive Weeks-Chandler-Andersen (WCA) potential.

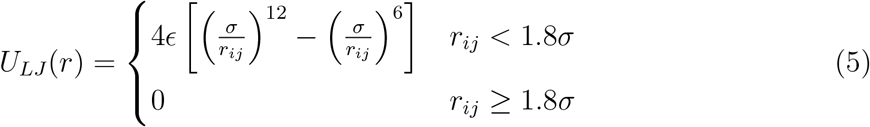

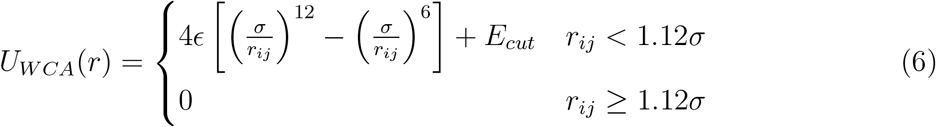

where *r*_*ij*_ is the distance between the *i*^*th*^ and *j*^*th*^ beads in 3D space, and *ϵ* determines the strength of the attractive interaction. *E*_*cut*_ is defined as 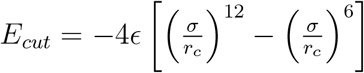, with *r*_*c*_ = 1.12*σ*.

In the simulation, the interaction between lamin and chromatin is also modeled using the attractive LJ/WCA potential. Additionally, the system includes a fixed boundary representing the nuclear membrane, which interacts attractively with lamin but is completely repulsive to chromatin. Another fixed boundary is purely repulsive to both chromatin and lamin. Table I summarizes the interaction parameters for different bead types.

**TABLE I:**
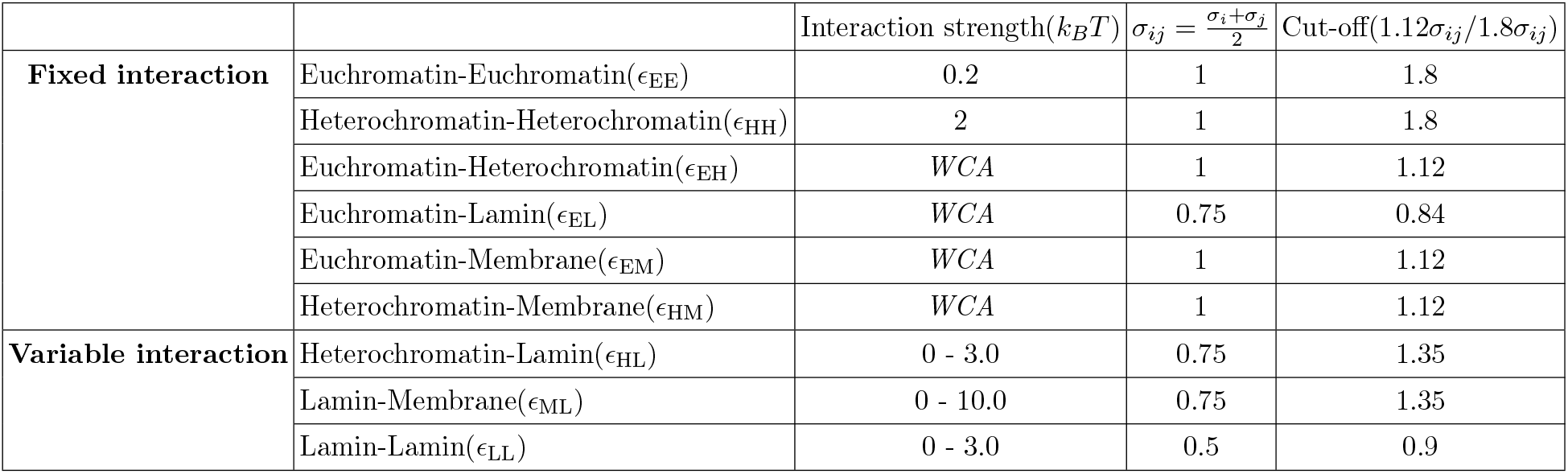
Interaction parameters betwen different components

| | | Interaction strength($k_B T$) | $\sigma_{ij} = \frac{\sigma_i + \sigma_j}{2}$ | Cut-off($1.12\sigma_{ij}/1.8\sigma_{ij}$) |
| --- | --- | --- | --- | --- |
| **Fixed interaction** | Euchromatin-Euchromatin($\epsilon_{EE}$) | 0.2 | 1 | 1.8 |
| | Heterochromatin-Heterochromatin($\epsilon_{HH}$) | 2 | 1 | 1.8 |
| | Euchromatin-Heterochromatin($\epsilon_{EH}$) | *WCA* | 1 | 1.12 |
| | Euchromatin-Lamin($\epsilon_{EL}$) | *WCA* | 0.75 | 0.84 |
| | Euchromatin-Membrane($\epsilon_{EM}$) | *WCA* | 1 | 1.12 |
| | Heterochromatin-Membrane($\epsilon_{HM}$) | *WCA* | 1 | 1.12 |
| **Variable interaction** | Heterochromatin-Lamin($\epsilon_{HL}$) | 0 - 3.0 | 0.75 | 1.35 |
| | Lamin-Membrane($\epsilon_{ML}$) | 0 - 10.0 | 0.75 | 1.35 |
| | Lamin-Lamin($\epsilon_{LL}$) | 0 - 3.0 | 0.5 | 0.9 |

**TABLE II:**
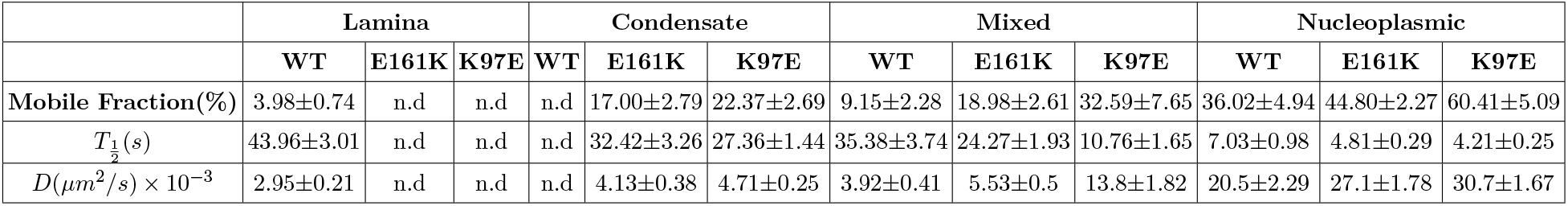
Mobile fractions, 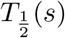, and diffusion coefficients obtained from the FRAP analysis for the different ROIs. Errors represents standad deviations.

|  | Lamina |  |  | Condensate |  |  | Mixed |  |  | Nucleoplasmic |  |  |
| --- | --- | --- | --- | --- | --- | --- | --- | --- | --- | --- | --- | --- |
|  | WT | E161K | K97E | WT | E161K | K97E | WT | E161K | K97E | WT | E161K | K97E |
| **Mobile Fraction(%)** | $3.98\pm0.74$ | n.d | n.d | n.d | $17.00\pm2.79$ | $22.37\pm2.69$ | $9.15\pm2.28$ | $18.98\pm2.61$ | $32.59\pm7.65$ | $36.02\pm4.94$ | $44.80\pm2.27$ | $60.41\pm5.09$ |
| $T_{\frac{1}{2}}(s)$ | $43.96\pm3.01$ | n.d | n.d | n.d | $32.42\pm3.26$ | $27.36\pm1.44$ | $35.38\pm3.74$ | $24.27\pm1.93$ | $10.76\pm1.65$ | $7.03\pm0.98$ | $4.81\pm0.29$ | $4.21\pm0.25$ |
| $D(\mu\text{m}^2/s) \times 10^{-3}$ | $2.95\pm0.21$ | n.d | n.d | n.d | $4.13\pm0.38$ | $4.71\pm0.25$ | $3.92\pm0.41$ | $5.53\pm0.5$ | $13.8\pm1.82$ | $20.5\pm2.29$ | $27.1\pm1.78$ | $30.7\pm1.67$ |

#### Simulation scheme

To investigate the organization of the lamin-chromatin composite system, we employed Langevin dynamics simulations, where the motion of each chromatin and lamin bead is described by the following equation:

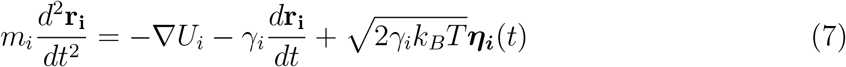

In this equation, *r*_*i*_ denotes the position of the *i*^*th*^ chromatin or lamin bead, *m*_*i*_ is its mass, and *U*_*i*_ represents the total interaction potential affecting the *i*_*th*_ bead. The friction coefficient *γ*_*i*_ determines the diffusion constant 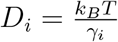 for each bead. We set *m*_*i*_ = 1 and *γ*_*i*_ = 1.0 for all beads. The last term in the equation accounts for random collisions due to solvent particles. The LAMMPS [104] software was utilized to numerically integrate the equations of motion using the standard velocity-Verlet algorithm, maintaining a constant temperature of *T* = 1.0 in reduced units. To ensure computational efficiency and numerical stability, we chose an integration time step of Δ*t* = 0.01*τ*_*Br*_, where *τ*_*Br*_ is the Brownian time, defined as the typical time for a bead to diffuse a distance comparable to its size (i.e.,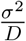, with *D* being the diffusion coefficient).

The initial configuration of the polymer was constructed as a self-avoiding polymer (SAW). Subsequently, 2000 lamin beads were randomly positioned within the simulation space. Equilibrated configurations were recorded after 4 *×* 10^7^ simulation steps for further analysis.

### H. Characterisation of simulation configurations

We examined several observables to quantify the system’s configuration within the parameter space of varying interaction strengths. Given the finite size and complexity of the system, there were broad crossovers between different classes of configurations as the interaction parameters were varied. In order to identify these crossover zones we identified macroscopic quantifications to distinguish the configurations, as discussed below. Specifically, we gathered 100 independent equilibrated configurations, each originating from different initial setups, for further analysis.

#### Lamin-lamin and membrane-lamin interaction plane

For *ϵ*_LL_ = 0.2, we characterised the different classes of configurations obtained by varying the energy scales *ϵ*_ML_ and *ϵ*_HL_. To quantify lamin presence near the membrane, we measured the number of lamin beads within 2.5*σ* of the membrane. A threshold of 300 lamin particles was used to distinguish configurations where lamin was concentrated near the membrane from those where it was absent. Next, we focused on identifying configurations where lamin beads were present within HC clusters versus those where the HC cluster cores lacked lamin. Using a cluster detection algorithm, we identified HC clusters and calculated the fraction of lamin beads within the radius of gyration of these clusters. We applied a threshold value of 0.3 to distinguish between configurations with lamin in HC clusters and those without. HC clusters tend to accumulate near the lamin-coated membrane due to the interaction parameter *ϵ*_HL_. To quantify this, we counted the number of HC beads located within 4.5*σ* of the membrane, with a threshold of 30 beads to determine whether HC clusters were present near the membrane. Furthermore, when the interaction between HC and lamin is sufficiently strong, HC clusters lose their sphericity and form uniform layers beneath the lamin. We calculated the radius of gyration (R_*g*_) of the HC clusters to differentiate between a compact HC cluster and an extended HC layer near the membrane. The boundary lines as shown in Fig. 2C were obtained by a smoothed representation of these different crossovers as derived from the quantifications.

#### Lamin-lamin and heterochromatin-lamin interaction plane

When laminmembrane interactions are weak (*ϵ*_ML_=3.0), or when these interactions are completely absent (*ϵ*_LM_=WCA), we characterised the configurations obtained in the interaction space of parameters *ϵ*_LL_ and *ϵ*_HL_. First, we performed a cluster analysis on lamin particles, defining a cluster as present when the average contained more than 20 lamin beads. Based on the presence or absence of such clusters, we identified regions of the interaction plane where lamin aggregates did not form. Next, we determined the proximity of clusters to the membrane by checking whether their center of mass was within 4.5*σ* of the membrane. We then differentiated this region from areas where lamin clusters formed both near the membrane and in the bulk nucleoplasm. We also investigated the lamin density within HC clusters, identifying a threshold value of *ϵ*_HL_ for each *ϵ*_LL_, above which lamin and HC clusters began to co-localize. Our analysis revealed that the highest lamin density occurred when HC clusters formed a core containing a substantial lamin aggregate, with HC coating the outside. Conversely, the lowest lamin density was observed when HC penetrated the lamin core, leading to a mixed lamin-HC cluster. An intermediate state was also identified, where partial mixing of HC within lamin clusters was observed. The boundary lines as shown in Fig. 2F,I were obtained by a smoothed representation of these different crossovers as derived from the quantifications.

## I. Three-dimensional fluorescent in situ hybridization (3D FISH)

Transfected Hela cells were given a brief wash with PBS and 5 minutes in ice-cold CSK buffer(0.1 M NaCl, 0.3 M sucrose, 3 mM MgCl2, 10 mM PIPES (pH 7.4), 1mM EGTA, 0.5 % Triton X-100). Cells were then fixed with 4% w/v PFA in room temperature for 10 minutes and washed with 0.1 M TrisHCl (pH 7.4), followed by 3 washes with PBS for 5 minutes each in room temperature. Cells were then incubated in 0.5% v/v Triton X100/PBS for 10 minutes followed by an incubation in 20% v/v Glycerol/PBS overnight, in 4°C. This incubation was followed by a freeze-thaw cycle of dipping the coverslips into liquid nitrogen for 30 seconds and thawing on a piece of tissue paper until the frozen layer disappeared, then immediately putting the coverslip back into 20% v/v glycerol. This cycle was repeated four times. Then the coverslips were washed in PBS for 10 minutes for 3 times followed by an incubation in 0.1 N HCl for 5 minutes in room temperature. Cells were then incubated twice for 3 minutes in 2X Saline Sodium Citrate (SSC) buffer followed by 3 washes in PBS, 5 minutes each. The coverslips with cells were then kept immersed in 50% v/v formamide (pH 7.0)/ 2X SSC solution for 24 hours at 4°C. Hybridization was performed with the red fluorochrome tagged chromosome13 probe (Applied Spectral Imaging, Israel). For each sample, 5 *µ*l probewas initially equilibrated for 5 minutes and denatured for 5 minutes at 80°C followed by an incubation at 37°C for 30 minutes. Then the probe was spotted on a clean glass slide and coverslips were mounted with cells facing the probes and sealed by nail-polish followed by a co-denaturation, performed at 80°C for 5 minutes. After denaturation the slides were kept in a humid chamber at 37°C for 3 days in the dark for hybridization. Post hybridization the coverslips were carefully withdrawn from top of the slide and washed with 50% v/v formamide (pH 7.0)/ 2X SSC solution at 45°C, thrice for 5 minutes each. This step was followed by 3 washes for 5 minutes each in 0.1X SSC buffer at 60°Cwith gentle agitation. Coverslips were then briefly rinsed with 0.1% v/v Tween-20/4X SSC. Finally, coverslips were washed in 2X SSC buffer and mounted with Vectashield Plus (Fisher Scientific, USA), sealed, and stored for image acquisition.

## J. Fluorescence recovery after Photobleaching (FRAP)

Following 24-hour of transfection, the cells were washed with PBS and transferred to CO_2_-independent media (Gibco, USA) following which the cells were equilibrated for 30 min at 37°C. All photobleaching experiments were conducted in NIKON TiE inverted microscope using λ405nm laser at 20 mW. The cells were maintained within a fully incubated chamber (Tokai Hit, Japan) at 37°C while being imaged with a 100X objective with a numerical aperture of 1.40, using a λ 488nm laser line with minimum laser power. A square region of interest (ROI) measuring 0.25 *µm*^2^ was designated for bleaching-cum-time-based fluorescent measurement. Three distinct categories were selected for bleaching experiments:

i. ROI containing nucleoplasmic fraction, while meticulously avoiding nuclear lamina or lamin A aggregates.
ii. ROI exclusively comprising wild type nuclear lamina or lamin A mutant aggregates, meticulously excluding the nucleoplasmic fraction.
iii. ROI encompassing approximately 50% of lamina/aggregates and the remaining nucleoplasmic fraction.

All ROIs were imaged for 5 seconds to obtain initial fluorescent intensity before undergoing bleaching. Bleaching of the ROIs was accomplished for 5 seconds followed by post-bleach recovery for 110 seconds post-bleaching at a shutter speed of 3.741 fps, utilizing 5% of 13 mW 488nm laser power with an active perfect focusing system. Any nuclear movement was eliminated by aligning the frames with respect to the first frame using the ND Analysis-Alignment module. The recovery of fluorescence was quantified using NIS-Elements software (Nikon) employing the ‘Time measurement’ function. 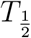 of the fluorescent recovery was calculated using FRAP module embedded in NIS-Elements by choosing the appropriate bleach point and recovery periods. For each time point, the data were normalized using the formula:

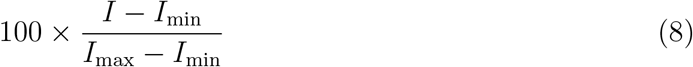

Mobile fraction was calculated using the formula:

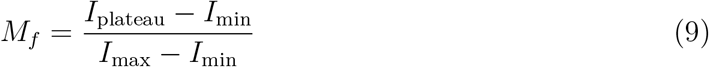

Where *I*_max_ represent maximum intensity (pre-bleach), *I*_plateau_ represent intensity after recovery, and *I*_min_ represent minimum intensity (post-bleach) Diffusion constant was calculated using the formula:

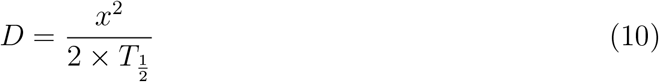

Where *x*^2^ represent bleach area. the detailed values obtained from each analysis (Mobile fractions, half-life of recovery 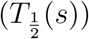, and diffusion coefficients) are mentioned in Table II.

## K. Statistical analysis

All the statistical analysis were done using GraphPad prism (Ver 8.4.2). Unpaired student t-test was used to determine the significance of the results with a 5% tolerance.

## V. AUTHOR CONTRIBUTIONS

K.S. and M.K.M. conceived and designed the project. S.D. developed the codes, performed simulations and analyzed the experimental images with input from all authors. S.N., S.D.S. and D.S. performed experiments, processed experimental images and analyzed data with input from all authors. S.D., S.N., D.S. and S.D.S. wrote the initial draft of the paper. M.K.M. and K.S. edited and prepared the final version of the draft.

## VI. ACKNOWLEDGEMENT

S.N., S.D.S. and D.S. acknowledges DAE, Govt. of India for the fellowship. S.D. acknowledges fellowship support from the PMRF, MoE, India. M.K.M acknowledges funding from Core Research Grant by Science and Engineering Research Board, India (Grant number: CRG/2022/008142). KSG acknowledges intramural project-RS4002 of DAE, Govt. of India.

